# Eulertigs: minimum plain text representation of *k*-mer sets without repetitions in linear time

**DOI:** 10.1101/2022.05.17.492399

**Authors:** Sebastian Schmidt, Jarno N. Alanko

**Affiliations:** University of Helsinki, Finland

**Keywords:** Spectrum preserving string sets, Eulerian cycle, Suffix tree, Bidirected arc-centric de Bruijn graph, k-mer based methods

## Abstract

A fundamental operation in computational genomics is to reduce the input sequences to their constituent *k*-mers. For maximum performance of downstream applications it is important to store the *k*-mers in small space, while keeping the representation easy and efficient to use (i.e. without *k*-mer repetitions and in plain text). Recently, heuristics were presented to compute a near-minimum such representation. We present an algorithm to compute a minimum representation in optimal (linear) time and use it to evaluate the existing heuristics. For that, we present a formalisation of arc-centric bidirected de Bruijn graphs and carefully prove that it accurately models the *k*-mer spectrum of the input. Our algorithm first constructs the de Bruijn graph in linear time in the length of the input strings (for a fixed-size alphabet). Then it uses a Eulerian-cycle-based algorithm to compute the minimum representation, in time linear in the size of the output.

**2012 ACM Subject Classification:** Applied computing → Computational biology; Theory of computation → Data compression; Theory of computation → Graph algorithms analysis; Theory of computation → Data structures design and analysis

## 1 Introduction

### Motivation

A *k*-mer is a DNA string of length *k* that is considered equal to itself and its reverse complement. A common pattern in bioinformatics is to reduce a set of input strings to their constituent *k*-mers. Such representations are at the core of many bioinformatics pipelines – see e.g. Schmidt et al. [23] or Brinda et al. [6] for an overview of applications. The wide-spread use of *k*-mer sets has prompted the question of what is the smallest *plain text representation* for a set of *k*-mers. Here, a plain text representation means a set of strings that have the same set of *k*-mers as the input strings, i.e. the *spectrum* is preserved. Such representations are also called *spectrum preserving string sets* (SPSS) [22], or simplitigs [6]. This has the following advantages over encoded representations:

■ When storing *k*-mer sets to disk, plain text may remove the need of decompression before usage, as some tools that usually take unitigs as input can take any other plain text representation without modification (e.g. Bifrost [13]).
■ Within an application, an encoded representation would require decoding whenever a *k*-mer is accessed, which may slow down the application a lot compared to when each *k*-mer is in RAM in plain text.

Further, in applications, it might be useful if the representation contains each *k*-mer exactly once. This is because some applications, like e.g. SSHash [21], are able to take any set of *k*-mers as input, but cannot easily deal with duplicate *k*-mers in the input.

### Related work

There are two heuristic approaches to the construction of a small SPSS without repetitions, namely *prophasm* [6] and *UST* [22]. While neither of these computes a minimum representation, Rahman et al. [22] also present a lower bound to the minimum size of any representation without repetition, and they show that they are within 3% of this lower bound in practice. They also present a counter-example showing that their lower bound is not tight. Small SPSSs without repetitions are used e.g. in SSHash [21] and are also computed by state-of-the-art de Bruijn graph compactors like Cuttlefish 2 [15].

When *k*-mer repetitions are allowed in an SPSS, there is a known polynomially computable minimum representation, namely *matchtigs* [23]. While matchtigs are expensive to compute, the authors also present a more efficient greedy heuristic that is able to compute a near-minimum representation on a modern server with no significant penalty in runtime (when compared to computing just unitigs), but a significant increase in RAM usage.

In [6, 23] the authors also showed that decreasing the size of an SPSS results in significantly better performance in downstream applications, i.e. when further compressing the representation with general purpose compressors, or when performing *k*-mer-based queries.

The authors of both [6] and [22] consider whether computing a minimum representation without repetitions may be NP-hard, as it is equivalent to computing a minimum path cover in a de Bruijn graph, which is NP-hard in general graphs by reduction from Hamiltonian cycle. However, computing a Hamiltonian cycle in a de Bruijn graph is actually polynomial [14]. The authors of [14] argue that de Bruijn graphs are a subclass of *adjoint* graphs, in which solving the Hamiltonian cycle problem is equivalent to solving the Eulerian cycle problem in the *original* of the adjoint graph, which can be computed in linear time^2^. However, the argument is only made for normal directed (and not bidirected) graphs, and thus is not applicable to our setup, where a *k*-mer is also considered equal to its reverse complement.

### Our contributions

Our first technical contribution is to carefully define the notion of a bidirected de Bruijn graph such that the spectrum of the input is accurately modelled in the allowed walks of the graph. Our definition also takes into account *k*-mers that are their own reverse complement. This technicality is often neglected in the literature, and sidestepped by requiring that the value of *k* is odd, in which case this special case does not occur. We give a suffix-tree-based deterministic linear-time algorithm to construct such a graph, filling a theory gap in the literature, as existing approaches [8, 15, 13, 1] depend on the value of *k* and/or are probabilistic due to the of use hashing, minimizers or Bloom filters, or do not use the reverse-complement-aware definition of *k*-mers [7].

Given the bidirected de Bruijn graph, we present an algorithm that computes a minimum plain text representation of *k*-mer sets without repetitions, which runs in output sensitive linear time. Steps 1 to 3 run in linear time in the number of nodes and arcs in the graph. In short, it works as follows:

1. Add breaking arcs into this graph to make it Eulerian.
2. Compute a Eulerian cycle in the resulting graph.
3. Break that cycle at the breaking arcs.
4. Output the strings spelled by the resulting walks.

The algorithm is essentially an adaption of the matchtigs algorithm [23], removing the possibility of joining walks by repeating *k*-mers. We give detailed descriptions for all these steps and prove their correctess in our bidirected de Bruijn graph model. Together with our linear-time de Bruijn graph construction algorithm, we obtain the main result of our paper:

#### Theorem 1.

*Let k be a positive integer and let I be a set of strings of length at least k over some alphabet* Σ. *Then we can compute a set of strings I’ of length at least k with minimum cumulative length and* CS_*k*_(*I*) = CS*_k_*(*I*’) *in O*(||*I*||log |Σ|) *time*.

where CS_*k*_(*I*) = CS_*k*_(*I*’) means that *I*’ is an SPSS of *I*, and ||*I*|| is the cumulative length of I (see Section 2 for accurate definitions). This gives a positive answer to the open question if a minimum SPSS without repetitions can be computed in polynomial time. Additionally, we give an easily computable tight lower bound on the size of a minimum SPSS without repetitions.

For our experiments, we have implemented steps 1 to 4 in Rust, taking the de Bruijn graph as given. The implementation is available on github: https://github.com/algbio/matchtigs. Our experimental evaluation shows that our algorithm does not result in significant practical improvements, but for the first time allows to benchmark the quality the heuristics prophasm and UST against an optimal solution. It turns out that both produce close-to-optimal results, but with a different distribution of computational resources.

Our work also shows that using arc-centric de Bruijn graphs can aid the intuition for certain problems, as in this case, the node-centric variant hides the relationship between Eulerian cycles and minimum SPSS without repetition.

### Organisation of the paper

In Section 2 we give preliminary definitions of well-known concepts. In Section 3 we define de Bruijn graphs and prove the soundness of the definitions. In Section 4 we show how to construct de Bruijn graphs by our definitions in linear time. In Section 5 we show how to construct a minimum SPSS without repetitions in linear time if the de Bruijn graph is given. In Section 6 we compare our algorithm and Eulertigs against strings computed with prophasm and UST on practical data sets.

## 2 Preliminaries

In this section we give the prerequisite knowledge required for this paper.

### 2.1 Bidirected graphs

In this section we define our notion of the bidirected graphs and the incidence model.

A multiset is defined as a set *M*, and an implicit function 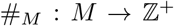 mapping elements to their multiplicities. The cardinality is defined as 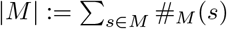.

An *alphabet* Σ is an ordered set, and an *Σ-word* is a string of characters of that set. String concatenation is written as *ab* for two strings *a* and *b*. The set Σ^*k*^ is the set of all Σ-words of length *k* and the set Σ* is the set of all Σ-words, including the empty word *ϵ*. Given a positive integer *k*, the *k-suffix* suf_*k*_(*w*) (*k-prefix* pre_*k*_(*w*)) of a word *w* is the substring of its last (first) *k* characters. A *k*-mer is a word of length *k*. A *complement function* over Σ is a function comp: Σ → Σ mapping characters to characters that is self-inverse (i.e. comp(comp(*x*)) = *x*). A *reverse complement* function for alphabet Σ is a function rc: Σ* → Σ* defined as rc((*w*_1_, …, *w_ℓ_*)):= (comp(*w_ℓ_*),…, comp(*w*_1_)), for some arbitrary complement function comp. On sets, rc is defined to compute the reverse complement of each element in the set. Note that rc is self-inverse. A *canonical k*-mer is a *k*-mer that is lexicographically smaller than or equal to its reverse complement.

Given an integer *k* and an alphabet Σ, the *k-spectrum* of a set of strings 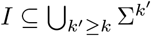 is a set of strings 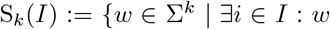 is substring of *i* or rc(*i*)}. The *canonical k-spectrum* of *I* is CS_*k*_(*I*):= {*w* ∈ S_*k*_(*I*) | *w* is canonical}. For simplicity, the spectrum and canonical spectrum are defined for a single string *w* as if it were a set {*w*}. A *spectrum preserving string set* of a set of strings *I* is a set of strings *I*’ such that CS_*k*_ (*I*) = CS_*k*_ (*I*’). The cumulative length of *I* is ||*I*||:=∑_*w I*_ |*w*|.

Our definition of a bidirected graph is mostly standard like in e.g. [17], however we allow self-complemental nodes that occur in bidirected de Bruijn graphs. A *bidirected graph* is a tuple *G* = (*V, E, c*) with a set of normal and *self-complemental* nodes *v ∈ V*, a set of arcs *e ∈ E,* and a function *c*: *V* → {1, 0} marking self-complemental nodes with 1, and normal nodes with 0. An *incidence* is a pair *vd,* where 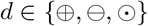 is called its *sign* (e.g. *v*⊕). The negation of a sign is defined as 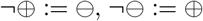. and 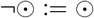. For self-complemental nodes *v* ∈ *V*, only incidences *v*⊙ are allowed, and for normal nodes only incidences *v*⊕ and 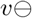 are allowed. An 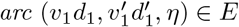 is a tuple of incidences and a unique identifier *η*, where *η* can be of any type. The *reversal* of an arc is denoted by 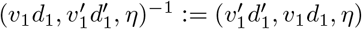. If not required, we may drop the identifier (i.e. just write 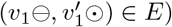. If a node *v* ∈ *V* is present with a ⊕ 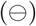 sign in an arc, then the arc is *outgoing (incoming)* from (to) *v*.

Note that, other than in standard directed graphs, in bidirected graphs arcs can be outgoing or incoming on both ends, and the order of the incidences in the arc does not affect if it is outgoing or incoming to a node. Further, our notation differs from that of standard bidirected graphs in that arcs have a direction. This is required because we will work with arc-centric de Bruijn graphs (see Section 3), which have labels on the arcs and not the nodes. Using the sign of the incidence pairs, it is possible to decide if a node is traversed forwards or backwards, but not if the arc is traversed forwards or backwards. But to decide which label (forwards or reverse complement) to use when computing the string spelled by an arc, the direction is relevant. See Figure 1 (a) for an example of a bigraph, which has labels that make it a de Bruijn graph as well.

**Figure 1.**
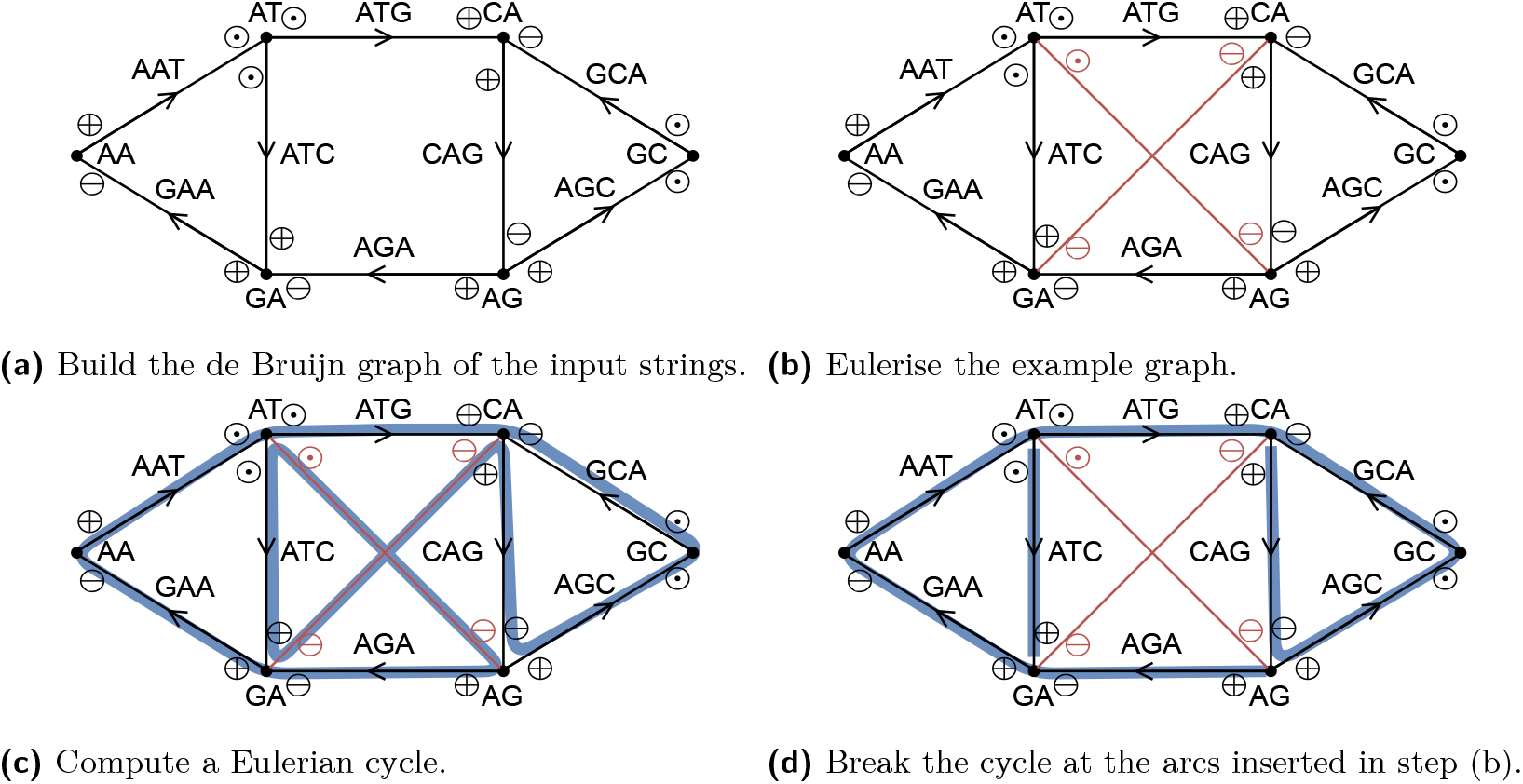
Overview of our algorithm executed on the input strings *{GAATG, ATCTGCT}* with *k* = 3. After step (d), the resulting spelled SPSS is *{ATC, AGAATGCTG}*.

A *walk* in a bigraph is a sequence of arcs 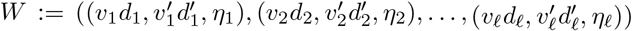 where for every *i* it holds that 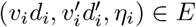 or 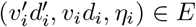 (we can arbitrarily walk over arcs forwards and reverse), and for every *i* < *ℓ* it holds that 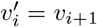 and 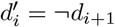. The *length* of a walk is *ℓ* = |*W*|. If 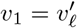 and 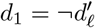, then *W* is a *cycle.* A bigraph is *connected,* if for each pair of nodes *v*_1_,*v*_2_ ∈ *V* there is a walk from *v*_1_ to *v*_2_.

For a node *v* ∈ *V*, the *multiset of incidences* is defined as 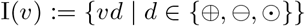, with multiplicities 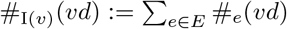 (treating the arcs as multisets such that selfloops count as two separate incidences). For a node *v* ∈ *V* that is not self-complemental, the *outdegree* is defined as 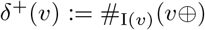, and the *indegree* is defined as 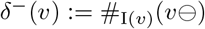. For a self-complemental node *v* ∈ *V*, the *degree* is defined as *δ*(*v*):= #_*I*(*v*)_(*v*⊙).

We define the *imbalance* of a node *v* ∈ *V* that is not self-complemental as the difference of its outdegree and indegree imbalance(*v*):= *δ*^+^(*v*) – *δ*^-^(*v*). For a self-complemental node *v* ∈ *V* the imbalance is defined as imbalance(*v*):= 1 if *δ*(*v*) is odd, and imbalance(*v*):= 0 otherwise. A node *v ∈ V* is called *unbalanced,* if imbalance(*v*) ≠ 0, and *balanced* otherwise.

A *labelled graph* is a bidirected graph *G* = (*V, E, c*) where the identifiers of arcs are strings over some alphabet Σ (e.g. 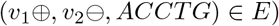).

### 2.2 Suffix arrays and suffix trees

Section 4 requires knowledge of suffix arrays and suffix trees. We assume the reader is familiar with these data structures, and briefly give the relevant definitions and properties below. We point the reader to Gusfield [12] and Mäkinen [18] for an in-depth treatment of the topics.

A suffix array *SA_T_* for a string *T* is an array of length |*T*| such that *SA_T_*[*i*] is the starting position of the lexicographically *i*-th suffix of *T*. The suffix array interval of a string *x* is the maximal interval [*i..j*] such that all the suffixes pointed by *SA_T_*[*i*],…, *SA_T_*[*j*] have *x* as a prefix, or the empty interval if *x* is not a substring of *T*.

A suffix tree of a string *T* is a compacted version of the trie of all suffixes of *T*, such that non-branching paths are merged into single arcs, with arcs pointing away from the root. The compactification concatenates the labels of the arcs on the compacted path. The nodes that were compacted away and are now in the middle of an arc are called implicit nodes, and the rest of the nodes are explicit. A *locus* (plural *loci*) is a node that is either explicit or implicit. A locus *v* is represented by a pair (*u, d*), where *u* is the explicit suffix tree node at the end of the arc containing *v* (*u* is equal to *v* if *v* is explicit), and *d* is the depth of locus *v* in the trie of loci. The suffix array interval of a node is the interval of leaves in the subtree of the node. The suffix array interval of an implicit locus (*u,d*) is the same as the suffix array interval of *u*.

The suffix tree can be constructed in linear time in |*T*| using e.g. Ukkonen’s algorithm [24]. The tree comes with a function child that takes an explicit node and a character, and returns the child at the end of the arc from that node whose label starts with the given character (if such node exists). This can be implemented in *O*(log |Σ|) time by binary searching over child pointers sorted by labels. The child function can also be easily implemented for implicit loci. Ukkonen’s algorithm also produces *suffix links* for the explicit nodes, which map from the suffix tree node of a string *cx* to the suffix tree node of string *x*. It is possible to emulate suffix links on the implicit loci using constant-time weighted level-ancestor queries [4] by mapping (*u, d*) ↦ (*f*_*d*-1_(*SL*(*u*)), *d* – 1), where *SL*(*u*) is the destination of a suffix link from *u*, and *f*_*d*-1_(*SL*(*u*)) is the furthest suffix tree ancestor from *SL*(*u*) at depth at least *d* – 1 in the trie of loci. The inverse pointers of suffix links are called *Weiner links*, and they can also be simulated on the implicit loci by mapping (*u, d*) ↦ (*WL*(*u, c*), *d* + 1), where *WL*(*u, c*) is the destination of a Weiner link from *u* with character *c*.

## 3 De Bruijn graphs

The *de Bruijn graph* of order *k* of a set of input strings *I* is defined as a labelled graph constructed by Algorithm 1. See Figure 1 (a) for an example. A de Bruijn graph computed by this algorithm has the following property (see Appendix B for some of the proofs of this section).

### Lemma 2

(Sound labels). *Let k be a positive integer and let I be a set of strings of length at least k. Let G =* (*V, E, c*) *be the de Bruijn graph of order k constructed from I. For all pairs of arcs* 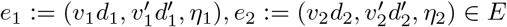 *it holds that*:

a. *(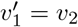 and 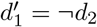) if and only if* suf_*k*-1_(*η*_1_) = pre_*k*-1_(*η*_2_),
b. *(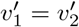 and 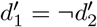) if and only if* suf_*k*-1_(*η*_1_) = pre_*k*-1_(rc(*η*_2_)),
c. *(v*_1_ = *v*_2_ *and* 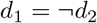 *if and only if* suf_*k*-1_(rc(*η*_1_)) = pre_*k*-1_(*η*_2_), *and*
d. (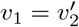 *and* 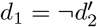*) if and only if* suf_*k*-1_(rc(*η*_1_)) = pre_*k*-1_(rc(*η*_2_)).

For a walk 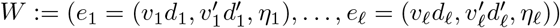 in a de Bruijn graph, its *sequence of k-mers* is *K*:= (*κ*_1_,…, *κ_ℓ_*), where for each *i* we define *κ_i_* as *η_i_* if *e_i_* ∈ *E*, and as rc(*η_i_*) if 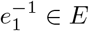. The string spell(*W*) is the string *spelled* by *W*, which is defined as its *collapsed* sequence of kmers, i.e. its sequence of *k*-mers gets concatenated while overlapping consecutive *k*-mers by *k* – 1. This is computed by Algorithm 2. We prove the following lemmas to show that our definition of the spell(·) function is sound for our purposes, i.e. correctly spells the string belonging to a walk in a de Bruijn graph.

### Algorithm 1 DeBruijnGraph

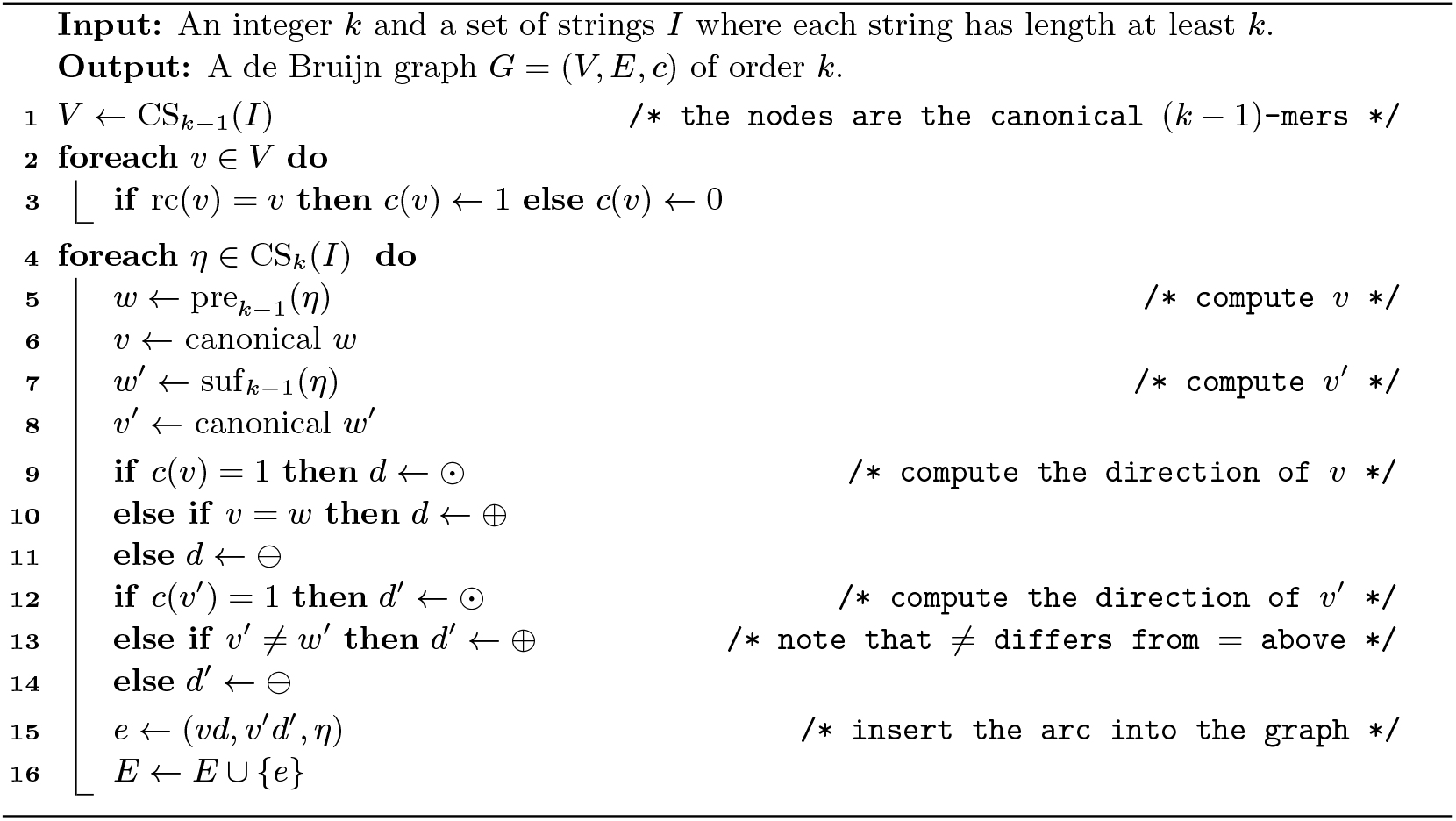

### Lemma 3

(Sound sequence of *k*-mers). *Let k be a positive integer and let I be a set of strings of length at least k. Let G* = (*V, E, c*) *be the de Bruijn graph of order k constructed from I. Let* 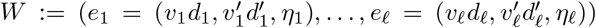 *be a walk in G, and K*:= (*κ*_1_,…, *κ_ℓ_*) *its sequence of k-mers. Then for each consecutive pair of kmers κ_i_*, *κ*_*i*+1_ *it holds that* suf_*k*-1_(*κ_i_*) = pre_*k*-1_(*κ*_*i*+1_).

We define the *sequence of k-mers K* = (*κ*_1_,…, *κ_ℓ_*) of a string *w* = (*a*_1_,…, *a*_*ℓ*+*k*-1_) by *κ_i_*:= (*a_i_*,…, *a*_*i*+*k*-1_) for each *i*.

#### Algorithm 2 Spell

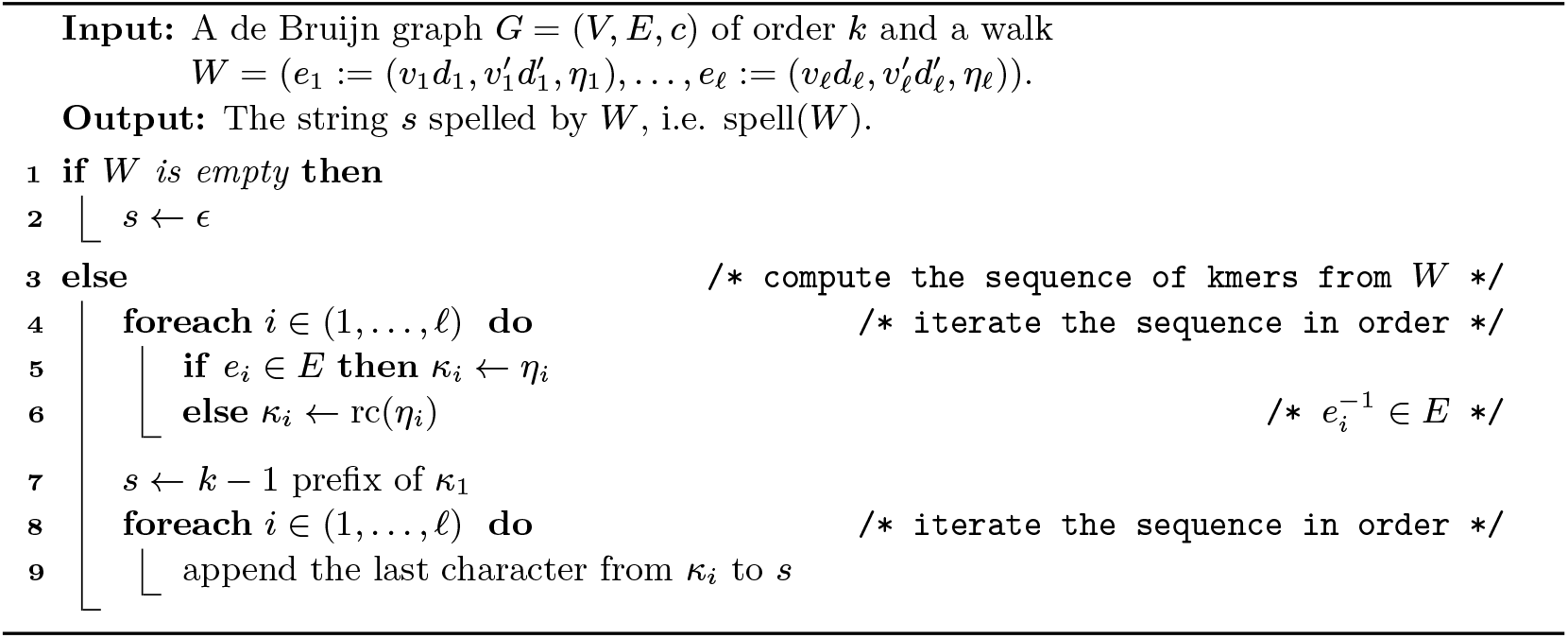

### Lemma 4

(Sound spell). *Let k be a positive integer and let I be a set of strings of length at least k. Let G* = (*V, E, c*) *be the de Bruijn graph of order k constructed from I. Let W be a walk in G, K_w_ its sequence of k-mers and* 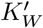 *the sequence of k-mers of* spell(*W*). *Then* 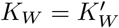.

### Lemma 5

(Complete representation). *Let k be a positive integer and let I be a set of strings of length at least k. Let G* = (*V, E, c*) *be the de Bruijn graph of order k constructed from I. Let w be a string with* 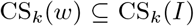. *Then there exists a walk W in G with* spell(*W*) = *w*.

**Proof.** Let *K_w_* = (*κ*_1_,…, *κ_ℓ_*) be the sequence of *k*-mers of *w*. We construct 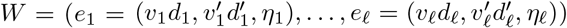 as follows: for each *i*, let *η_i_* be the canonical of *κ_i_* and *f_i_* ∈ *E* be the arc whose identifier is *η_i_*. We set *e_i_* = *f_i_* if *κ_i_* is canonical, and 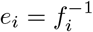 otherwise.

For *W* to fulfil the definition of a walk we need that 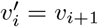 and 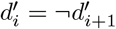 for all *i*. Using Lemma 2, we get:

■ If *e_i_*,*e*_*i*+1_ ∈ *E*, then suf_*k*-1_(*η_i_*) = suf_*k*-1_(*κ_i_*) = pre_*k*-1_(*κ*_*i*+1_) = pre_*k*-1_(*η*_*i*+1_). Therefore, by Lemma 2 (a), it holds that 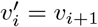 and 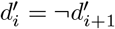.
■ If 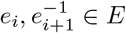, then suf_*k*-1_(*η_i_*) = suf_*k*-1_ (*κ_i_*) = pre_*k*-1_ (*κ*_*i*+1_) = pre_*k*-1_(rc(*η*_*i*+1_)). Therefore, by Lemma 2 (b), it holds that 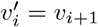 and 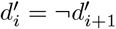.
■ If 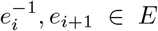, then suf_*k*-1_(rc(*η_i_*)) = suf_*k*-1_(*κ_i_*) = pre_*k*-1_(*κ*_*i*+1_) = pre_*k*-1_(*η*_*i*+1_). Therefore, by Lemma 2 (c), it holds that 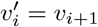 and 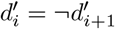.
■ If 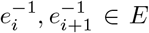, then suf_*k*-1_(rc(*η_i_*)) = suf_*k*-1_(*κ_i_*) = pre_*k*-1_(*κ*_*i*+1_) = pre_*k*-1_(rc(*η*_*i*+1_)). Therefore, by Lemma 2 (d), it holds that 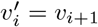 and 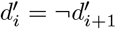.

To complete the proof we need to show that spell(*W*) = *w*. By definition, the sequence of *k*-mers *K_W_* of *W* is equivalent to *K_W_*. And since *W* is a walk, by Lemma 4 we get that the sequence of *k*-mers of spell(*W*) is equivalent to *K_W_*, and therefore spell(*W*) = *w*.

A *walk cover* 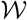 of a bigraph *G* is a set of walks such that for each arc *e ∈ E* it holds that *e* is part of some walk 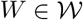, or *e*^-1^ is part of some walk 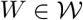.

### Theorem 6

(Dualism between SPSS and walk cover). *Let k be a positive integer and let I and I’ be sets of strings of length at least k. Let G* = (*V, E, c*) *be the de Bruijn graph of order k constructed from I. Then it holds that* CS_*k*_(*I*) = CS_*k*_(*I*’), *if and only if there is a walk cover* 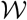 *in G that spells the strings in I*′.

**Proof.** If 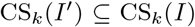, then for each string *w*’ ∈ *I*’ it holds that 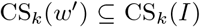. Therefore, by Lemma 5, there exists a walk *w* in *G* with spell(*w*) = *w*′. Then, the set of all such walks 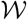 spells *I*’. Further, because 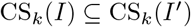, the identifier *η* of each arc *e ∈ E* is in CS_*k*_(*I*’), and therefore in the sequence of kmers *K_w’_* of some string *w*’ ∈ *I*’ (possibly as a reverse complement). By Lemma 4 it holds that *K_w’_* = *K_w_*, where *K_w_* is the sequence of *k*-mers of walk *w*. By the definition of the sequence of *k*-mers of a walk, this implies that *w* visits *e* (possible in reverse direction). Since this holds for each *e ∈ E*, it holds that 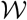 is a walk cover of *G*.

Assume that there is a walk cover 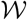 in *G* that spells the strings in *I*′, and let 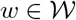 be a walk, *K_w_* its sequence of *k*-mers, *w*’:= spell(*w*) and *K_w’_* the sequence of *k*-mers of *w*’. Then, by Lemma 4, *K_w_* = *K_w’_*, which, by the definition of the sequence of *k*-mers of a walk implies that 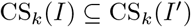. And since 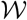 is a walk cover of *G*, we get CS_*k*_(*I*) = CS_*k*_(*I*’).

### Corollary 7.

*By setting I* = *I’ in Theorem 6 we can confirm that our definition of a de Bruijn graph is sound in that there is a set of walks that spells the strings used for its construction*.

A *compacted* de Bruijn graph is constructed from a de Bruijn graph by contracting all nodes *v ∈ V* that are either self-complemental and have exactly two arcs that have exactly one incidence to *v* each, or that are not self-complemental and have exactly one incoming and one outgoing arc. For simplicity, we use uncompacted de Bruijn graphs in our theoretical sections, however all results equally apply to compacted de Bruijn graphs.

## 4 Linear-time construction of compacted bidirected de Bruijn graphs

In this section, we fill a gap in the literature by describing on a high level an algorithm to construct the bidirectional de Bruijn graph of a set of input strings in time linear in the total length of the input strings, independent of the value of *k*.

### 4.1 Algorithm

Let *I* = {*w*_1_,… *w_m_*} be the set of input strings. Consider the following concatenation:

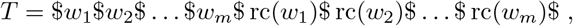

where $ is a special character outside of the alphabet Σ of the input strings. We require an index on *T* that can answer the following queries: `extendRight, extendLeft, contractRight` and `contractLeft` in constant time. The extension operations take as input a character *c* ∈ Σ and the interval of a string *x* in the suffix array of *T*, and return the suffix array intervals of *xc* in the case of `extendRight` and *cx* in the case of `extendLeft`. The contraction operations are the inverse operations of these, mapping the suffix array intervals of *xc* to *x* in the case of `contractRight` and *cx* to *x* in the case of `contractLeft`. For efficiency, we also require operations `enumerateRight` and `enumerateLeft`, which take a string *x* and give all characters such that `extendRight` and `extendLeft` respectively return a non-empty interval, in time that is linear in the number of such characters. Implementations for all the six subroutines are given in Section 4.2.

Using these operations, we can simulate the regular non-bidirected de Bruijn graph of *T*. Each *k*-mer of the input strings for a fixed *k* corresponds to a disjoint interval in the suffix array of *T*. The nodes are represented by their suffix array intervals. The outgoing arcs from a (*k* – 1)-mer *x* are those characters *c* where `extendRight`(*x, c*) returns a non-empty interval. We can enumerate all the characters *c* with this property in constant time using `enumerateRight`(*x*). The incoming arcs can be enumerated symmetrically with the `enumerateLeft`(*x*). Finally, we can find the destination or origin of an arc labelled with *x* by running a `contractLeft` or `contractRight` operation respectively on *x*.

To construct the bidirected de Bruijn graph, we merge together nodes that are the reverse complement of each other. To find which nodes are complemental, we scan the input strings *I* while maintaining the suffix array interval of the current *k*-mer using `extendRight` and `contractLeft` operations, while at the same time maintaining the suffix array interval of the reverse complement using `extendLeft` and `contractRight` operations. Whenever we merge two nodes, we combine the incoming and outgoing arcs, assigning the incidences of the arcs according to the incidence rules in our definition. We are able to tell in constant time which *k*-mer of a pair of complemental *k*-mers is canonical by comparing the suffix array intervals of the *k*-mers: the *k*-mer whose suffix array interval has a smaller starting point is the canonical *k*-mer. If the starting points are the same, the *k*-mer is self-complemental.

Using the `enumerateRight` and `enumerateLeft` functions, we can check if a node would be contracted in a compacted de Bruijn graph. By extending *k*-mers over such nodes, we can in linear time also output only the arcs and nodes of a compacted de Bruijn graph. For storing the labels, we use one pointer into the input strings to store a single *k*-mer, as well as a flag that is set whenever the label is not canonical. If a label has multiple *k*-mers, then we store the remaining *k*-mers as explicit strings, however without their overlap with the “pointer-*k*-mer”. This way, we can store each label in *O*(*ℓ*) space, where *ℓ* is the number of *k*-mers in the label. We additionally store the first and last character of each label, as an easy way to make the spell function run in output sensitive linear time.

### 4.2 Implementation of the subroutines

All required the subroutines `extendRight, extendLeft, contractRight, contractLeft, enumerateRight` and `enumerateLeft` can be implemented with the suffix tree of *T* by simulating the trie of the suffix tree loci as described in Section 2.2. The suffix array intervals of explicit nodes can be stored with the nodes, so that we can operate on loci (*u, d*) and retrieve the suffix array intervals on demand. The operation `extendRight` follows an arc from a locus to a child, and the operation `contractRight` is implemented by going to the parent of the current locus. The operation `contractLeft` follows a suffix link from the current locus, and `extendLeft` follows a Weiner link. The operations `enumerateRight` and `enumerateLeft` are implemented by storing the children and the Weiner links from explicit suffix tree nodes as neighbor lists. The total number of these links is linear in |*T*| [18]. With this implementation, the slowest operations are `extendRight` and `extendLeft`, taking *O*(log |Σ|) time to binary search the neighbor lists. We therefore obtain the following result:

#### Theorem 8.

*The compacted arc-centric bidirected de Bruijn graph of order k of a set of input strings I from the alphabet* Σ *can be constructed in time O*(||*I*|| log |Σ|).

We note that the same operations can also be implemented on top of the bidirectional BWT index of Belazzougui and Cunial [2], using the data structures of Belazzougui et al. [3] for the enumeration operations. This gives an index that supports all the required subroutines in *constant time.* The drawback of the bidirectional BWT index is that only randomized construction algorithms are known, but the expected time is still linear in |*T*|. We leave as an open problem the construction of the compacted arc-centric bidirected de Bruijn graph in deterministic linear time independent of the alphabet size.

## 5 Linear-time minimum SPSS without repetitions

Let *I* be a set of strings. To compute an SPSS without repetitions we first build a compacted de Bruijn graph *G* from *I*. Because of Theorem 6, finding an SPSS is equivalent to finding a walk cover in *G*. Further, with Lemma 4, we get that an SPSS without repetitions is equivalent to a walk cover that visits each arc exactly once (either once forwards, or once reverse, but not both forwards and reverse). We call such a walk cover a *unique walk cover.*

For minimality, observe that the cumulative length of an SPSS *S* relates to its equivalent set of walks 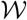 as follows:

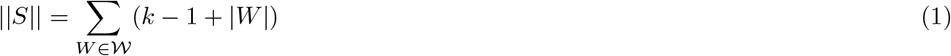

This is because in Algorithm 2, in Line 7, *k* – 1 characters are appended to the result, and then in the loop in Line 8, one additional character per arc in *W* is appended. We cannot alter the sum 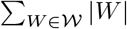, since we need to cover all arcs in *G*. However we can alter the number of strings, and decreasing or increasing this number by one will decrease or increase the cumulative length of *S* by *k* – 1. Therefore, finding a minimum SPSS of *I* without repetitions equals finding a unique walk cover of *G* that has a minimum number of walks.

Note that computing a minimum SPSS in a bigraph that is not connected is equivalent to separately computing an SPSS in each maximal connected subgraph. Therefore we restrict to connected bigraphs from here on.

### 5.1 A lower bound for an SPSS without repetitions

Using the imbalance of the nodes of a bigraph, we can derive a lower bound for the number of walks in a walk cover.

#### Lemma 9.

*Let v ∈ V be an unbalanced node in a bigraph G* = (*V, E, c*). *Then in a unique walk cover* 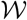 *of G, either at least* | imbalance(*v*)| *walks start in v, or at least* | imbalance(*v*)| *walks end in v*.

**Proof.** If *v* is self-complemental, then its imbalance is 1, so by definition *v* has an odd number of incident arcs. Each walk that does not start or end in *v* needs to enter and leave *v* via two distinct arcs whenever it visits *v*. But since the number of incident arcs is odd, there is at least one arc that cannot be covered this way, implying that a walk needs to start or end in this arc.

If *v* is not self-complemental and has a positive imbalance, then it has imbalance(*v*) more outgoing arcs then incoming arcs. Since walks need to leave *v* with the opposite sign than they entered *v*, at least imbalance(*v*) arcs cannot be covered by walks that do not start or end in *v*. If *v* has negative imbalance, the situation is symmetric.

#### Definition 10

(Imbalance of a bigraph). *The imbalance* imbalance(*G*) *of a bigraph G* = (*V, E, c*) *is the sum of the absolute imbalance of all nodes*∑| imbalance(*v*)|.

#### Theorem 11

(Lower bound). *Let G be a bigraph. A walk cover* 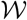 *of G has a minimum string count of* imbalance(*G*)/2.

**Proof.** Let *v ∈ V* be an unbalanced node. Then, by Lemma 9 at least | imbalance(*v*)| walks start in *v* or at least | imbalance(*v*)| walks end in *v*. Since each walk has exactly one start node and one end node, 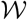 has a minimum string count of imbalance(*G*)/2.

### 5.2 Eulerising a bigraph

A directed graph is called *Eulerian*, if all nodes have indegree equal to outdegree, i.e. are balanced [10]. If the graph is strongly connected^3^, then this is equivalent to the graph admitting a *Eulerian cycle,* i.e. a cycle that visits each arc exactly once. The same notion can be used with bidirected graphs, using our definition of imbalance.

#### Definition 12

(Eulerian bigraph). *A bigraph is* Eulerian, *if all nodes have imbalance zero*. A connected bigraph can be transformed into a Eulerian bigraph by adding arcs using Algorithm 3. See Figure 1 (b) for an example.

#### Algorithm 3 Eulerise

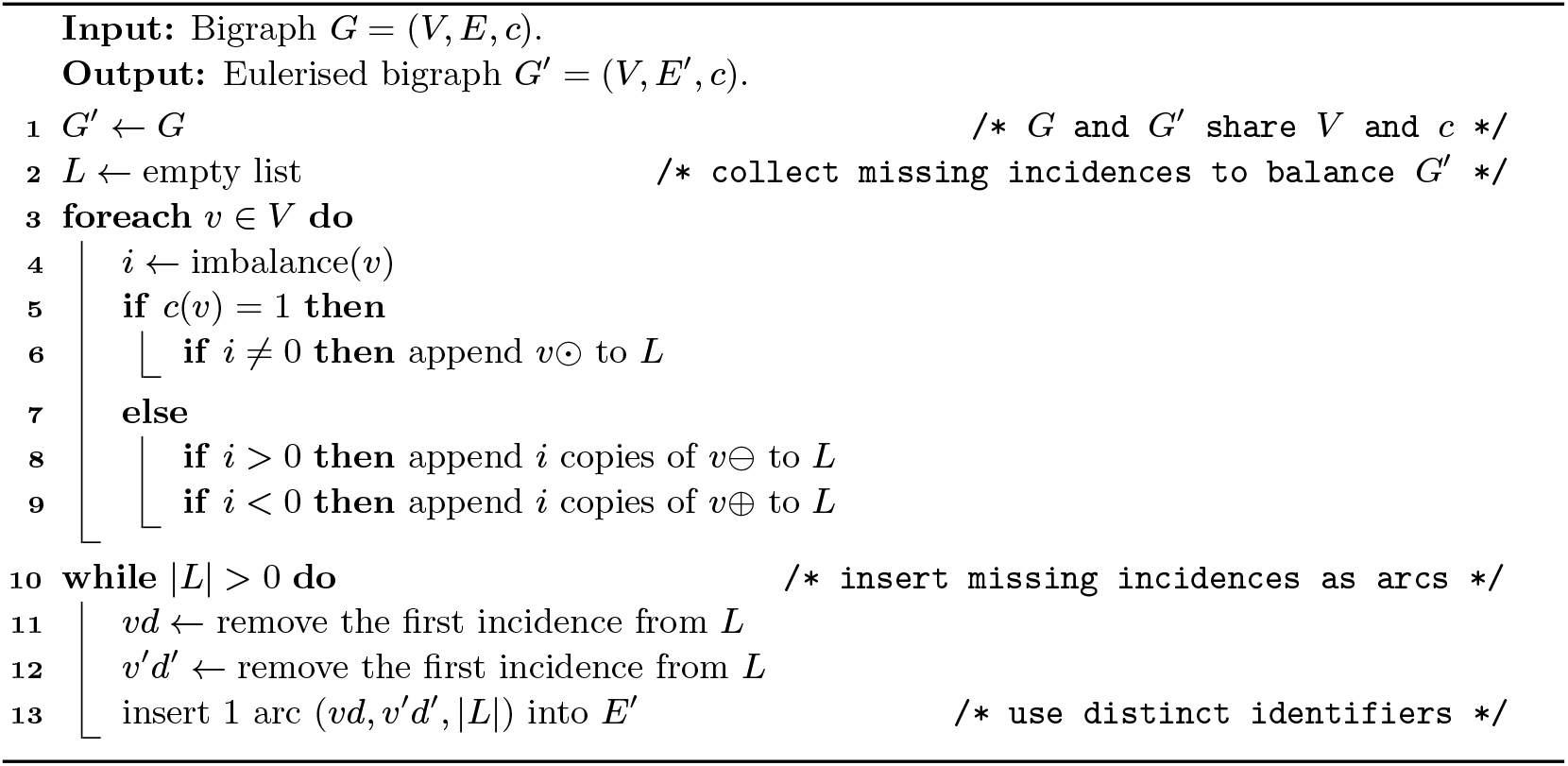

#### Lemma 13.

*The imbalance of a bigraph is even.*

**Proof.** Adding or removing an arc changes the imbalance of two nodes by 1, or of one node by two. In both cases, the imbalance of the graph can only change by –2, 0, or 2. Since the imbalance of a graph without arcs is 0, this implies that there can be no graph with odd imbalance.

#### Lemma 14.

*Given a connected bigraph G* = (*V,E,c*), *Algorithm 3 outputs a Eulerian bigraph G*’ = (*V, E’, c*).

**Proof.** Algorithm 3 is well-defined, since by Lemma 13, it holds that *L* has even length in each iteration of the loop in Line 10, so the removal operation in Line 12 always has something to remove.

The output of Algorithm 3 is a valid bigraph, since for self-complemental nodes *v* ∈ *V*, only incidences *v*⊙ are added to *G*’, and for not self-complemental nodes *v* ∈ *V*, only incidences *v*⊕ and 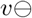 are added to *G*’.

Further, the output is a Eulerian bigraph, because for all *v* ∈ *V*, it holds that imbalance(*v*) is 0, by the following argument:

a. If *c*(*v*) = 1 and *v* has imbalance zero in *G*, then its imbalance stays the same in *G*’. If it has imbalance 1, then one incident arc is inserted, making its degree even and its imbalance therefore zero.
b. If *c*(*v*) = 0 and *v* has positive imbalance *i* in *G*, then *i* incoming arcs are added to *v* (counting incoming self-loops twice), and no outgoing arcs are added. Therefore, it has imbalance zero in *G*’. By symmetry, if *v* has negative imbalance in *G*, it has imbalance zero in *G*’.

#### Lemma 15.

*Given a bigraph G* = (*V,E,c*), Algorithm 3 terminates after O(|*V*| + |*E*|) *steps*.

**Proof.** For the list data structure we choose a doubly linked list, and for the graph an adjacency list (and array with an entry for each node containing a doubly linked list for the arcs).

The loop in Line 3 runs |*V*| times and each iteration runs in *O*(|imbalance(*v*)|) for a node *v*, because a doubly linked list supports appending in constant time. The sum of absolute imbalances of all nodes cannot exceed 2|*E*|, because each arc adds at most 1 to the absolute imbalance of at most two nodes, or adds at most 2 to the absolute imbalance of at most one node. Therefore, the length of list *L* after completing the loop is at most 2| *E*|, and the loop runs in *O*(|*V*| + |*E*|) time.

The loop in Line 10 runs at most |*L*| ≤ 2|*E*| times and performs only constant-time operations, since *L* is a doubly linked list and we can insert arcs into an adjacency list in constant time. Therefore, this loop also runs in *O*(|*V*| + |*E*|) time.

With Lemmas 14 and 15 we get the following.

#### Theorem 16.

*Algorithm 3 is correct and runs in O*(|*V*| + |*E*|) *time*.

### 5.3 Computing a Eulerian cycle in a bigraph

After Eulerising the bigraph, we can compute a Eulerian cycle using Algorithm 4. We do this similarly to Hierholzer’s classic algorithm for Eulerian cycles [10]. First we find an arbitrary cycle. Then, as long as there are unused arcs left, we search along the current cycle for unused arcs, and find additional cycles through such unused arcs. We integrate each of those additional cycles into the main cycle. See Figure 1 (c) for an example of a Eulerian cycle.

#### Algorithm 4 EulerianCycle

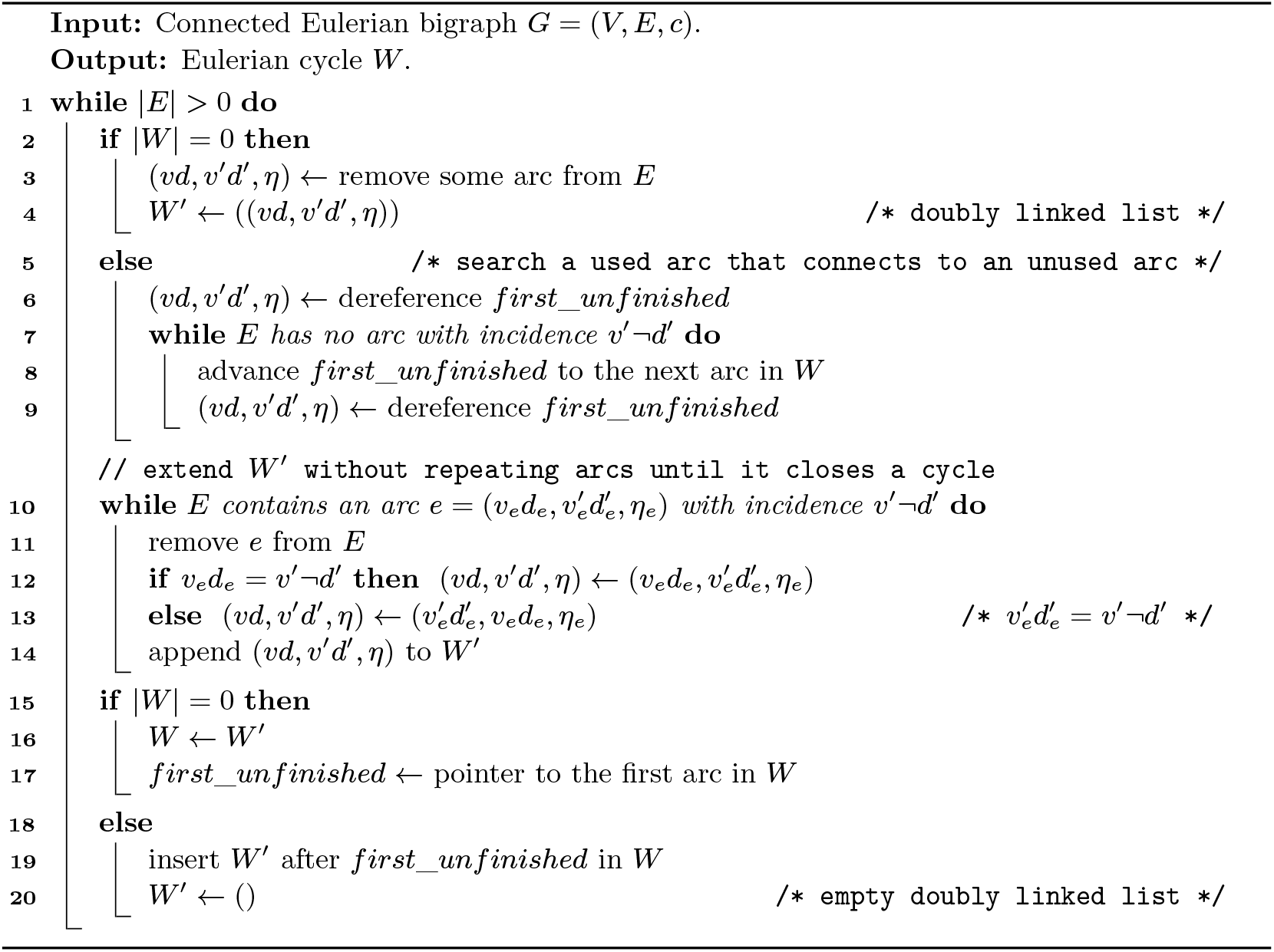

#### Lemma 17.

*Given a connected Eulerian bigraph G* = (*V,E,c*), *Algorithm 4 terminates and outputs a Eulerian cycle W*.

**Proof.** For 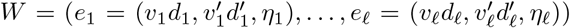 to be a Eulerian cycle, it must be a cycle that contains each arc exactly once.

The sequence *W*’ constructed by the loop in Line 10 is a walk by construction, and since *G* is Eulerian it is a cycle after the loop terminates. After finding the initial cycle in the first iteration of the outer loop, each additional cycle is started from a node on the initial cycle, and is a cycle again. Therefore it can be inserted into the original cycle without breaking its cycle property.

Since each arc is deleted when being added to *W*’, there is no duplicate arc in *W*. And if the algorithm terminates, then |*E*| = 0 (Line 1), so *W* contains all arcs.

For termination, consider that if *W* is not complete after the first iteration of the outer loop, then the loop in Line 7 searches for an unused arc using the *first_unfinished* pointer. Since the prefix of *W* up to including *first_unfinished* is never modified (Line 19), and *first_unfinished* is only advanced when its pointee cannot reach any arc anymore, it holds that no arc in *W* can reach an arc in *E* when *first_unfinished* gets advanced over the end of *W*. Since *G* was initially Eulerian and only Eulerian cycles have been removed from *G*, this implies that all nodes visited by *W* are still balanced and therefore have no incident arcs anymore. And since *G* was originally connected, *W* has visited all nodes, i.e. |*E*| = 0. Therefore, *first_unfinished* cannot be advanced over the end of *W*, because the outer loop terminates before that.

To complete the proof of termination, consider that in each iteration of the outer loop, at least one arc gets removed from *E*. In the first iteration, this happens at least in Line 3, and in all following iterations, this happens in Line 11.

#### Lemma 18.

*Given a connected Eulerian bigraph G* = (*V,E,c*), *Algorithm 4 terminates after O*(|*V*| + |*E*|) *steps*.

**Proof.** We use a doubly linked list for *W* and *W’*, and an adjacency list for *G*. Then all lines can be executed in constant time.

The loop in Line 10 removes one arc from *E* each iteration, so it runs at most |*E*| times in total (over all iterations of the outer loop). The loop in Line 7 advances *first_unfinished* each iteration. Since the algorithm is correct by Lemma 17, |*W*| ≤ |*E*| and *first_unfinished* never runs over the end of *first_unfinished,* so the loop runs at most |*E*| times in total (over all iterations of the outer loop).

The condition for the loop in Line 10 is true at least once in each iteration of the outer loop, since the preceding branch sets up (*vd, v’d’, η)* such that it has a successor (in the first iteration because of Eulerianess). So in each iteration of the outer loop, at least one arc gets removed, so the outer loop runs at most | *E*| times in total.

As a result, all loops individually run at most |*E*| times, therefore Algorithm 4 terminates after *O*(|*V*| + |*E*|) steps.

With Lemmas 17 and 18 we get the following.

#### Theorem 19.

*Algorithm 4 is correct and runs in O*(|*V*| + |*E*|) *time*.

### 5.4 Computing a minimum SPSS without repetitions

We convert the Eulerian cycle into a walk cover of the original bigraph by breaking it at all arcs inserted by Algorithm 3, and removing those arcs (see Figure 1 (d) for an example). This results in a walk cover with either one walk, if Algorithm 3 inserted zero or one arcs, or imbalance(*G*)*/*2 arcs, if Algorithm 3 inserted more arcs. By Theorem 11, this is a minimum number of walks, and therefore the SPSS spelled by these walks is minimum as well. Constructing the de Bruijn graph takes *O*(||*I*|| logΣ) time, and it has *O*(||*I*||) *k*-mers, so it holds that |*V*| ∈ *O*(||*I*||) and |*E*| ∈ *O*(||*I*||). Further, spelling the walk cover takes time linear to the cumulative length of the spelled strings. Since we compute a minimum representation, it holds that the output is not larger than the total length of the input strings. Therefore we get:

#### Theorem 1.

*Let k be a positive integer and let I be a set of strings of length at least k over some alphabet* Σ. *Then we can compute a set of strings I*′ *of length at least k with minimum cumulative length and* CS_*k*_(*I*) = CS_*k*_(*I*’) *in O*(||*I*|| log |Σ|) *time*.

## 6 Experiments

We ran our experiments on a server running Linux with two 64-core AMD EPYC 7H12 processors with 2 logical cores per physical core, 1.96TiB RAM and an SSD. Our data sets are the same as in [23], and we also adapted their metrics *cumulative length* (CL), which is the total count of characters in all strings, and *string count* (SC), which is the number of strings. Our implementation does not use the formalisation of bidirected graphs introduced in this work, but instead uses the formalisation from [23]. For constructing de Bruijn graphs, we do not implement our purely theoretical linear time algorithm, since practical de Bruijn graph construction is a well-researched field [8, 13, 15, 9, 20, 19], and we want to focus more on computing the compressed representation from unitigs. UST only supports unitigs constructed by BCALM2 [8], since it needs certain additional data. BCALM2 is not a linear time algorithm, but works efficient in practice. Therefore, we use BCALM2 to construct a node-centric de Bruijn graph, and then convert it to an arc-centric variant using a hash table.

Our experimental pipeline is constructed with [16] and using the bioconda software repository [11]. We ran all multithreaded tools with up to 28 threads and never used more than 128 cores of our machine at once to prevent hyperthreading from affecting our timing. The code to reproduce our experiments is available at https://doi.org/10.5281/zenodo.6538261.

The performance figures are all very similar, with two exceptions. Prophasm does not support parallel computation at the moment, therefore its runtime is much higher. Compared to that, all other algorithms use parallel computation to compute unitigs, but computing the final tigs from unitigs seems to be negligible compared to computing the de Bruijn topology. Moreover, running UST or Eulertigs on read data sets of larger genomes consumes significantly more memory than computing just unitigs. This is likely because BCALM2 uses external memory to compute unitigs, while the other tools simply load the whole set of unitigs into memory.

It is notable that the Eulertigs algorithm is always slower than UST. This may be because of the Eulertig algorithm being more complex, but also because our loading and storing routines might not be as efficient. While UST uses node-centric de Bruijn graphs and can therefore directly make use of the topology output by BCALM2 (which is a fasta file with arcs stored as custom annotations), we need to convert the graph into arc-centric format. This is supported by e.g. the B. mori short read data set, on which the computation of Eulertigs uses only 11% of the runtime for the algorithm itself, while 89% are from loading the graph (including the conversion to arc-centric) and storing the result.

In terms of CL, we see that the SPSS computed with UST mostly remains within the expected 3% of the lower bound, but they are up to 5% above the lower bound on more compressible data sets. The SPSS computed by prophasm is very close to the optimum in all cases, and we assume that this difference in quality is because prophasm extends paths both forwards and backwards, while the UST heuristic merely extends them forwards.

Looking at SC, we see that Eulertigs are always the lowest, which is due to the string count directly being connected to the cumulative length by Equation (1). This also explains the correlation between CL and SC, which can be observed in all cases.

## 7 Conclusions

We have presented a linear and hence optimal algorithm for computing a minimum SPSS without repetitions for a fixed alphabet size. This closes the open question about its complexity raised in [6, 22]. Using our optimal algorithm, we were able to accurately evaluate the existing heuristics and show that they are very close to the optimum in practice. Further, we have published our algorithm as a command-line tool on github, allowing it to easily be used in other projects.

Further, we have presented how bidirected de Bruijn graphs can be formalised without excluding any corner cases. We have also shown how such a graph can be constructed in linear time for a fixed-size alphabet. The construction of the compacted arc-centric bidirected de Bruijn graph in linear time independent of the alphabet size stays an open problem.

**Table 1.** Experiments on references and read sets of single genomes with *k* = 51 and a min abundance of 10 for human and 1 for the others. The CL and SC ratios are compared to the CL-optimal Eulertigs. For time and memory, we report the total time and maximum memory required to compute the tigs from the respective data set. BCALM2 directly computes unitigs, while UST- and Eulertigs require a run of BCALM2 first before they can be computed themselves. Prophasm can only be run for *k* ≤ 32, which does not make sense for large genomes. The number in parentheses behind time and memory indicates the slowdown/increase over computing just unitigs with BCALM2. BCALM2 was run with 28 threads, while all other tools support only one thread. The lengths of the genomes are 100Mbp for C. elegans, 482Mbp for B. mori and 3.21Gbp for H. sapiens and the read data sets have a coverage of 64x for C. elegans, 58x for B. mori and 300x for H. sapiens.

| genome | algorithm | CL ratio | SC ratio | time [s] | memory [GiB] |
| --- | --- | --- | --- | --- | --- |
| C. elegans (reads) | unitigs | 1.789 | 2.831 | 1888 | 5.97 |
|  | UST | 1.035 | 1.080 | 2738 (1.45) | 15.2 (2.54) |
|  | Eulertigs | 1 | 1 | 3735 (1.98) | 25.0 (4.19) |
| B. mori (reads) | unitigs | 1.912 | 3.136 | 7737 | 9.36 |
|  | UST | 1.050 | 1.118 | 10937 (1.41) | 52.4 (5.60) |
|  | Eulertigs | 1 | 1 | 13793 (1.78) | 79.4 (8.48) |
| H. sapiens (reads) | unitigs | 1.418 | 2.143 | 56966 | 13.0 |
|  | UST | 1.016 | 1.044 | 57736 (1.01) | 16.4 (1.26) |
|  | Eulertigs | 1 | 1 | 58861 (1.03) | 29.2 (2.25) |
| C. elegans | unitigs | 1.060 | 3.154 | 54.7 | 1.22 |
|  | UST | 1.002 | 1.089 | 58.0 (1.06) | 1.22 (1.00) |
|  | Eulertigs | 1 | 1 | 65.9 (1.21) | 1.22 (1.00) |
| B. mori | unitigs | 1.262 | 3.310 | 224 | 3.32 |
|  | UST | 1.018 | 1.156 | 258 (1.16) | 3.32 (1.00) |
|  | Eulertigs | 1 | 1 | 315 (1.41) | 3.32 (1.00) |
| H. sapiens | unitigs | 1.195 | 3.532 | 3166 | 10.0 |
|  | UST | 1.015 | 1.192 | 3369 (1.06) | 10.0 (1.00) |
|  | Eulertigs | 1 | 1 | 3717 (1.17) | 10.0 (1.00) |

**Table 2.**
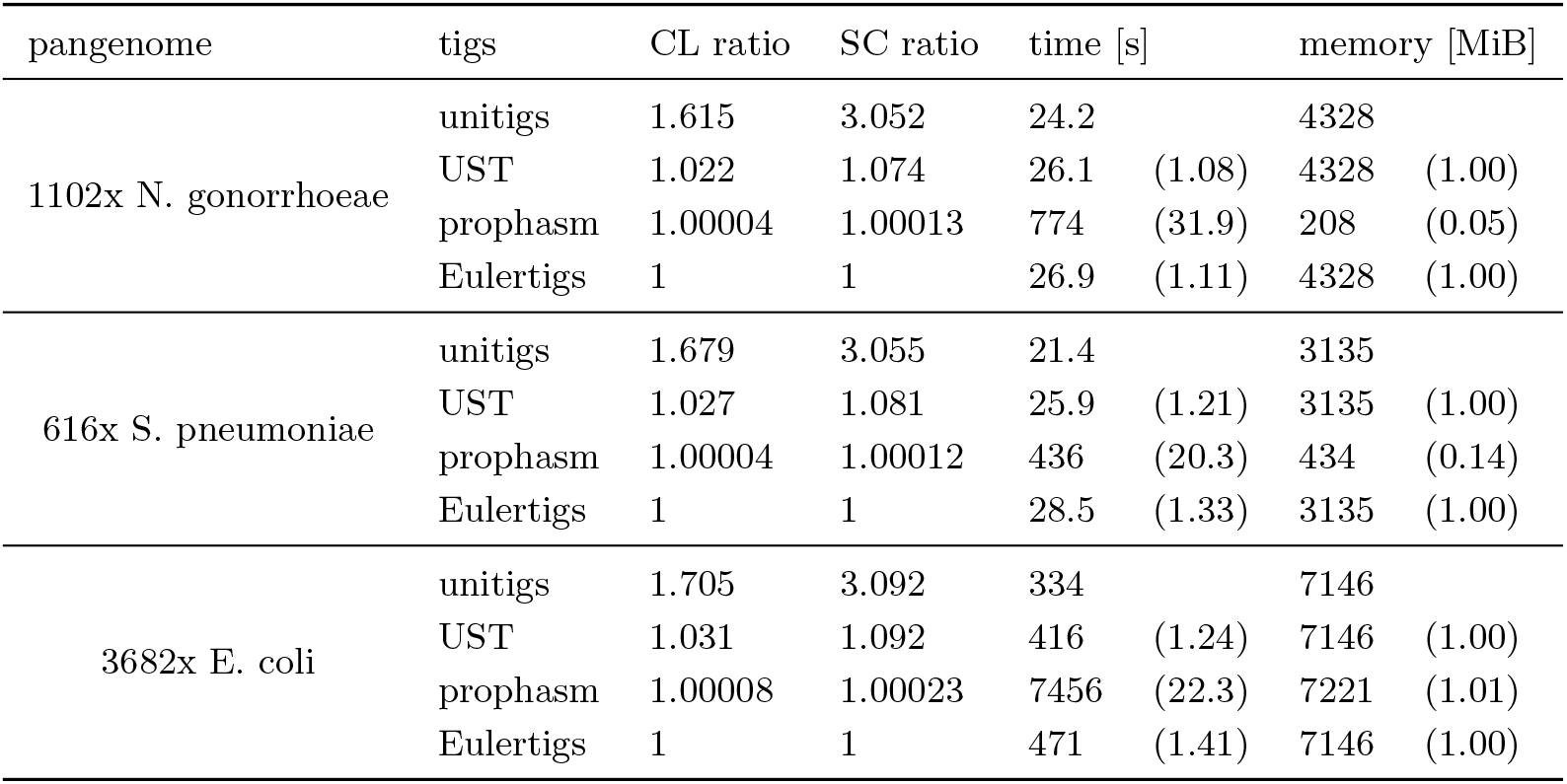
Experiments on (references of) pangenomes with *k* = 31 and a min abundance of 1. The CL and SC ratios are compared to the CL-optimal Eulertigs. For time and memory, we report the total time and maximum memory required to compute the tigs from the respective data set. BCALM2 directly computes unitigs, while UST- and Eulertigs require a run of BCALM2 first before they can be computed themselves. Prophasm is run directly on the source data. The number in parentheses behind time and memory indicates the slowdown/increase over computing just unitigs with BCALM2. BCALM2 was run with 28 threads, while all other tools support only one thread. The N. gonorrhoeae pangenome contains 8.36 million unique kmers, the S. pneumoniae pangenome contains 19.3 million unique kmers and the E. coli pangenome contains 341 million unique kmers.

| pangenome | tigs | CL ratio | SC ratio | time [s] | memory [MiB] |
| --- | --- | --- | --- | --- | --- |
| 1102x *N. gonorrhoeae* | unitigs | 1.615 | 3.052 | 24.2 | 4328 |
|  | UST | 1.022 | 1.074 | 26.1 (1.08) | 4328 (1.00) |
|  | prophasm | 1.00004 | 1.00013 | 774 (31.9) | 208 (0.05) |
|  | Eulertigs | 1 | 1 | 26.9 (1.11) | 4328 (1.00) |
| 616x *S. pneumoniae* | unitigs | 1.679 | 3.055 | 21.4 | 3135 |
|  | UST | 1.027 | 1.081 | 25.9 (1.21) | 3135 (1.00) |
|  | prophasm | 1.00004 | 1.00012 | 436 (20.3) | 434 (0.14) |
|  | Eulertigs | 1 | 1 | 28.5 (1.33) | 3135 (1.00) |
| 3682x *E. coli* | unitigs | 1.705 | 3.092 | 334 | 7146 |
|  | UST | 1.031 | 1.092 | 416 (1.24) | 7146 (1.00) |
|  | prophasm | 1.00008 | 1.00023 | 7456 (22.3) | 7221 (1.01) |
|  | Eulertigs | 1 | 1 | 471 (1.41) | 7146 (1.00) |

## A Authors contributions

JNA and SS discovered the problem, SS solved the problem when the de Bruijn graph is given and wrote most of the manuscript, JNA designed the linear-time de Bruijn graph construction algorithm and wrote Section 4. SS implemented the algorithm and conducted and evaluated the experiments. All authors reviewed and approved the final version of the manuscript.

## B Omitted proofs

### Lemma 2

(Sound labels). *Let k be a positive integer and let I be a set of strings of length at least k. Let G* = (*V, E, c*) *be the de Bruijn graph of order k constructed from I. For all pairs of arcs* 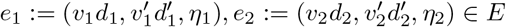 *it holds that*:

a. (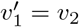 *and* 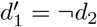) *if and only if* suf_*k*-1_(*η*_1_)=pre_*k*-1_(*η*_2_),
b. (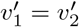 *and* 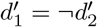*) if and only if* suf_*k*-1_ (*η*_1_) = pre_*k*-1_(rc(*η*_2_)),
c. (*v*_1_ = *v*_2_ *and* 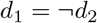*) if and only if* suf_*k*-1_ (rc(*η*_1_)) = pre_*k*-1_(*η*_2_), and
d. (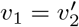 *and* 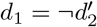*) if and only if* suf_*k*-1_ (rc(*η*_1_)) = pre_*k*-1_(rc(*η*_2_)).

**Proof.** Observe that the values of *w* and *w*′ computed in Lines 5 and 7 of Algorithm 1 are equal to pre_*k*-1_(*η*_1_) and suf_*k*-1_(*η*_1_) for *e*_1_ and equal to pre_*k*-1_(*η*_2_) and suf_*k*-1_(*η*_2_) for *e*_2_. Further, observe that the values of *v* and *v*’ computed in Lines 6 and 8 are equal to *v*_1_ and 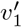 for *e*_1_ and equal to *v*_2_ and 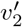 for *e*_2_. This makes *v*_1_, 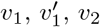, *v*_2_ and 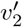 the canonicals of pre_*k*-1_(*η*_1_), suf_*k*-1_(*η*_1_), pre_*k*-1_(*η*_2_) and suf_*k*-1_(*η*_2_). Finally, observe that the sign values *d* and *d’* computed in Lines 9–14 are equal to *d*_1_ and 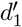 for *e*_1_ and equal to *d*_2_ and 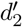 for *e*_2_.

a. If 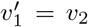 and 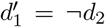, then 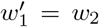 for all possible values of 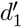, and therefore suf_*k*-1_ (*η*_1_) =pre_*k*-1_(*η*_2_). If suf_*k*-1_(*η*_1_) = pre_*k*-1_(*η*_2_), then 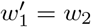, and therefore 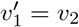 because 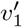 and *v*_2_ are the canonicals of 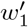 and *w*_2_. Additionally, 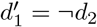 for all possible values of 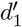.
b. If 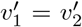 and 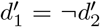, then 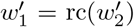 for all possible values of 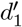, and therefore suf_*k*-1_ (*η*_1_) = rc(suf_*k*-1_ (*η*_2_)) = pre_*k*-1_(rc(*η*_2_)). If suf_*k*-1_(*η*_1_) = pre_*k*-1_(rc(*η*_2_)), then 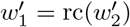, and therefore 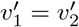 because 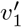 and 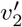 are the canonicals of 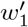 and 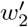. Additionally, 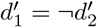 for all possible values of 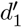.
c. If *v*_1_ = *v*_2_ and 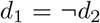, then rc(*w*_1_) = *w*_2_ for all possible values of *d*_1_, and therefore suf_*k*-1_(rc(*η*_1_)) = rc(pre_*k*-1_(*η*_1_)) = pre_*k*-1_(*η*_2_). If suf_*k*-1_(rc(*η*_1_)) = pre_*k*-1_(*η*_2_), then *w*_1_ = rc(*w*_2_), and therefore *v*_1_ = *v*_2_ because *v*_1_ and *v*_2_ are the canonicals of *w*_1_ and *w*_2_. Additionally, 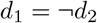 for all possible values of *d*_1_.
d. This case is equivalent to the first case when swapping *e*_1_ and *e*_2_, because 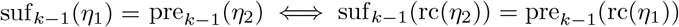.

### Lemma 3

(Sound sequence of *k*-mers). *Let k be a positive integer and let I be a set of strings of length at least k. Let G* = (*V, E, c*) *be the de Bruijn graph of order k constructed from I. Let* 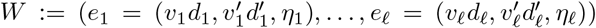 *be a walk in G, and K*:= (*κ*_1_,…, *κ_ℓ_*) *its sequence of k-mers. Then for each consecutive pair of kmers κ_i_, κ*_*i*+1_ *it holds that* suf_*k*-1_(*κ_i_*) = pre_*κ*-1_(*κ*_*i*+1_).

**Proof.** Let *i* ∈ {1,…, *ℓ* – 1}. By the definition of *walk* it holds that 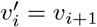 and 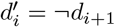. We can apply Lemma 2 case by case.

a. If 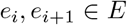, then by Lemma 2 (a), it holds that suf_*k*-1_(*η_i_*) equals pre_*k*-1_(*η*_*i*+1_). By definition, *κ_i_* = *η_i_* and *κ*_*i*+1_ = *η*_*i*+1_, so suf_*k*-1_(*κ_i_*) = pre_*k*-1_(*κ*_*i*+1_).
b. If 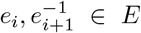, then by Lemma 2 (b) applied to 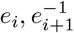, it holds that suf_*k*-1_(*η_i_*) equals pre_*k*-1_(rc(*η*_*i*+1_)). By definition, *κ_i_* = *η_i_* and *κ*_*i*+1_ = rc(*η*_*i*+1_), so suf_*k*-1_(*κ_i_*) = pre_*k*-1_(*κ*_*i*+1_)
c. If 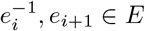, then by Lemma 2 (c) applied to 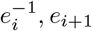, it holds that suf_*k*-1_(rc(*η_i_*)) equals pre_*k*-1_(*η*_*i*+1_). By definition, *κ_i_* = rc(*η_i_*) and *κ*_*i*+1_ = *η*_*i*+1_, so suf_*k*-1_(*κ_i_*) = pre_*k*-1_(*κ*_*i*+1_).
d. If 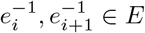, then by Lemma 2 (d) applied to 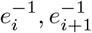, it holds that suf_*k*-1_(rc(*η_i_*)) equals pre_*k*-1_(rc(*η*_*i*+1_)). By definition, *κ_i_* = rc(*η_i_*) and *κ*_*i*+1_ = rc(*η*_*i*+1_), so suf_*k*-1_(*κ_i_*) = pre_*k*-1_(*κ*_*i*+1_).

### Lemma 4

(Sound spell). *Let k be a positive integer and let I be a set of strings of length at least k. Let G* = (*V, E, c*) *be the de Bruijn graph of order k constructed from I. Let W be a walk in G, K_W_ its sequence of k-mers and 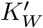 the sequence of k-mers of* spell(*W*). *Then* 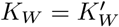.

**Proof.** Let (*κ*_1_,…, *κ_ℓ_*):= *K_W_*. We use induction over the length of *W*. For an empty *W*, *K* is empty, spell(*W*) is empty, and therefore *K*’ is empty as well. For |*W*| = 1, Algorithm 2 outputs spell(*W*) = *κ*_1_ and it holds that 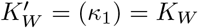.

For |*W*| ≥ 2 we consider that 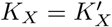 holds for a prefix *X* of *W* with |*X*| = |*W*| – 1. When *i* = |*W*| at the beginning of the loop in Line 8, then *s* = spell(*X*). By Lemma 3 it holds that the last *k* – 1 characters of *s* are equal to the first *k* – 1 characters of *κ_ℓ_*. Therefore, by appending the last character from *κ_ℓ_* to *s*, *κ_ℓ_* is appended to 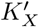 forming 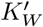. Therefore, last *k*-mer of 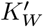 equals the last *k*-mer of *K_w_*, and the first *ℓ* – 1 *k*-mers of 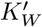 equal those of *K_W_* by induction.

## C Pseudocode for linear-time construction of compacted de Bruijn graphs

The pseudocode for computing a compacted de Bruijn graph in linear time is given by Algorithm 6 which uses Algorithm 5 as a subroutine. The data structure *D* used by the algorithms is that described in Section 4. Note that if we compute the arc labels as plain strings as in Algorithm 1, we need up to *O*(*k*) bits to store a single-*k*-mer arc. And since arcs are not substrings of input strings (but potentially combinations of input strings), we would need a string set of up to *O*(*k*||*I*||) characters to store all arc labels without referring to the input strings. This contradicts the algorithm being linear in ||*I*||. However, we can store the labels as tuples (*p, η, q, r*), where *pηq* is the label where *p* and *q* are explicit strings while *η* is a pointer to a *k*-mer in the input. If *r* is true, then the label must be reverse complemented to match that defined by Algorithm 1. With this fix, the size of each label is linear in the number of *k*-mers it represents, and in total the de Bruijn graph represents *O*(||*I*||) *k*-mers.

The comparison on Line 16 of Algorithm 6 can be done in linear time in |*η*_1_| + |*η*_2_ | by finding the suffix array intervals of *η*_1_*ηη*_2_ and rc(*η*_1_*ηη*_2_) with `extendLeft` and `extendRight` from *η* and rc(*η*) respectively, and comparing the starts of the intervals. This way, the total time taken by all those comparisons is proportional to the sum of |*η*_1_ | + |*η*_2_| over all unitigs, which is linear in ||*I*|| because each character of *η*_1_ and *η*_2_ can be mapped to a distinct edge in the non-compacted de Bruijn graph of ||*I*||. Therefore, the algorithm can be implemented to run in *O*(||*I*||) time.

Our pseudocode does not compute the first and last character of each arc-label, but this can be easily computed in constant time using *w_i_*, *η*_1_ and *η*_2_ in Algorithm 6.

### Algorithm 5 FindUnitigEnd

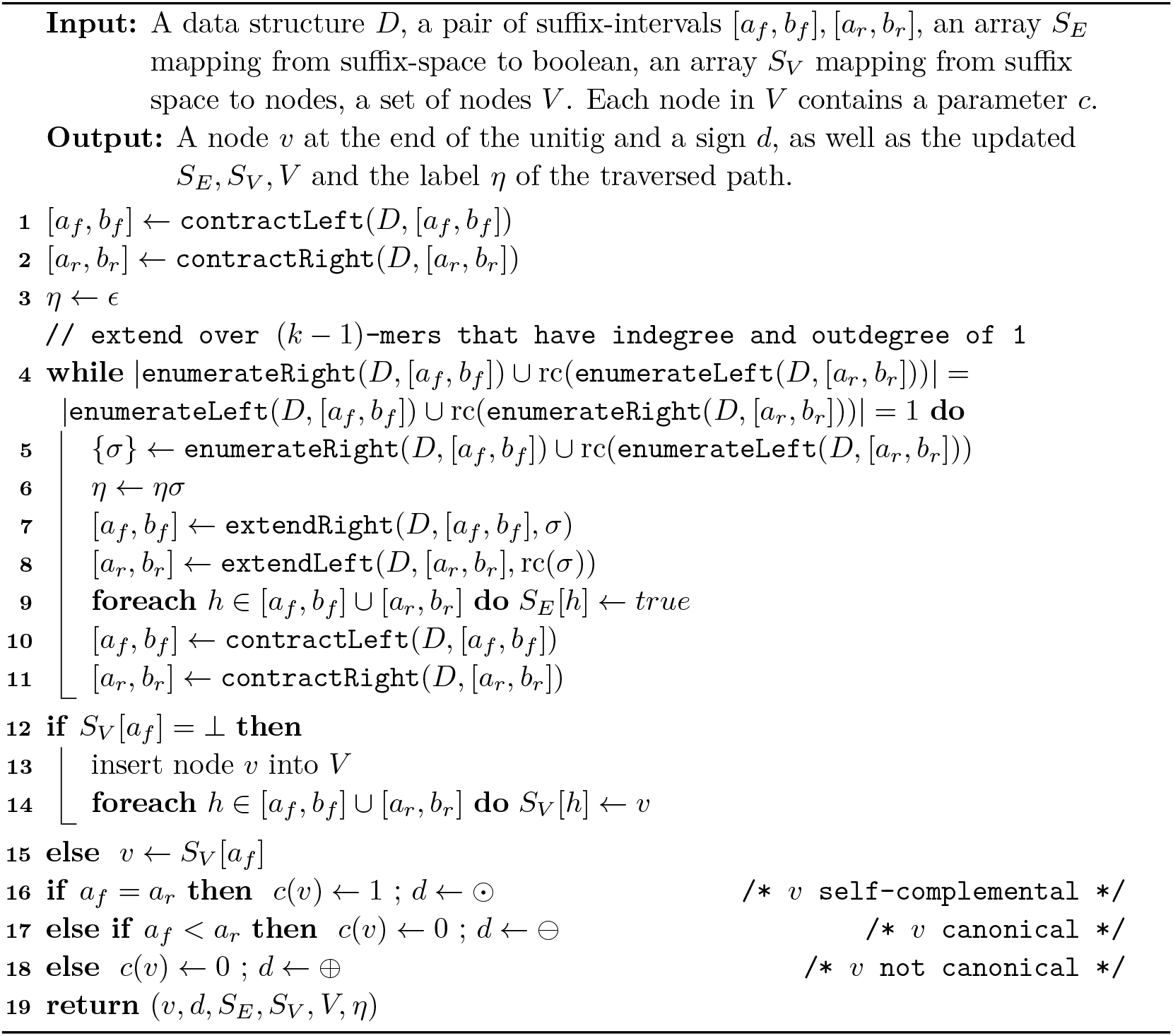

### Algorithm 6 LinearCompactedDbg

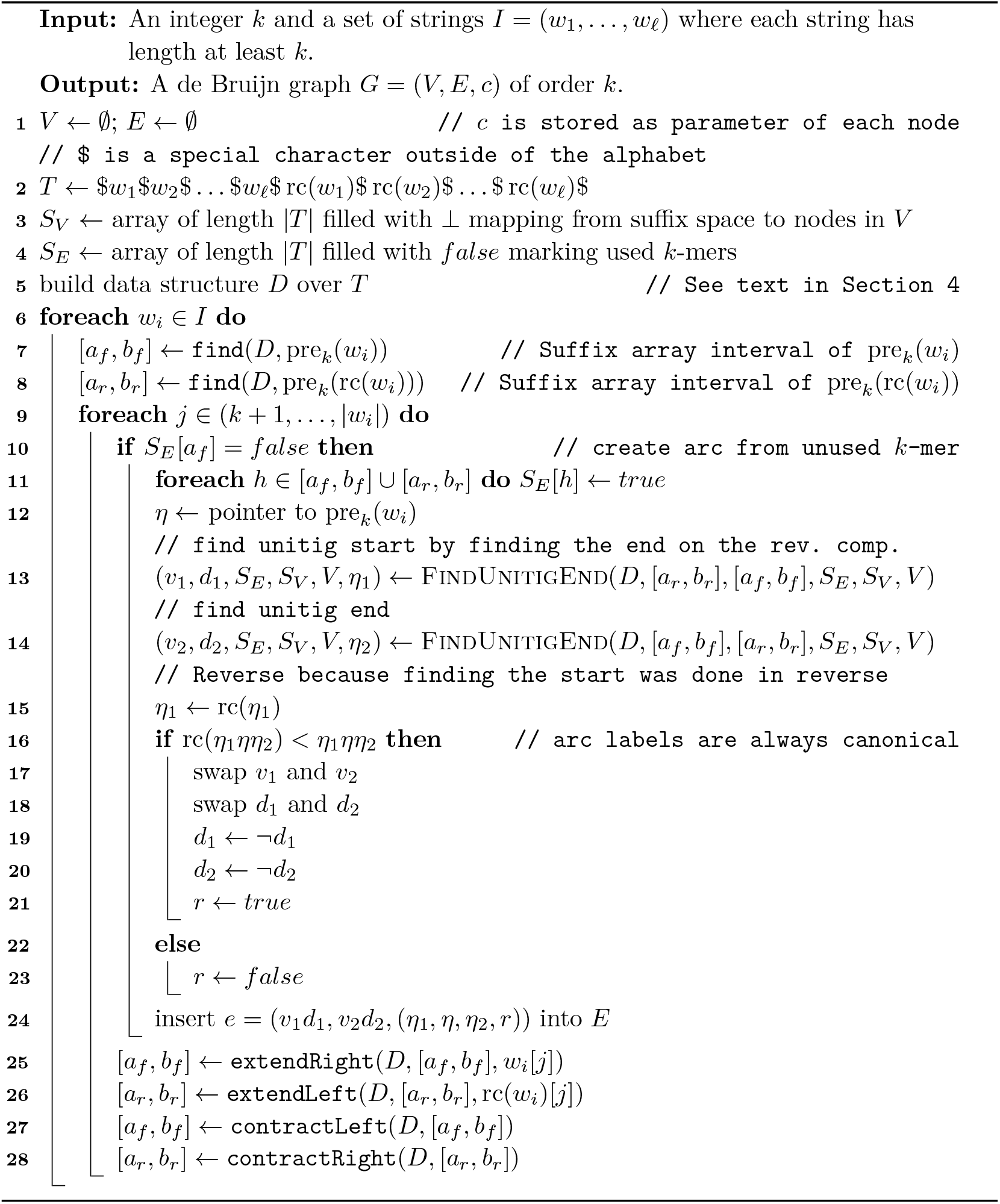

## Footnotes

2 The original of an adjoint graph can be computed by splitting each node *v* into two nodes *v*′ and *v*″ such that *v*′ keeps the incoming arcs, and *v*″ the outgoing arcs as in [5, Figure 4]. Then, the graph is a collection of complete bipartite graphs [5]. These graphs can be contracted into single nodes, and then we add an arc between the contracted representations of each *v*′ and *v*″. This can be computed in linear time and is the original graph, since all nodes have become arcs again, and the arcs have the correct predecessors and successors.

3 Strongly connected means that there is a directed path from each node *vi* to each node *v*_2_.

